# Regulation of STING activation by phosphoinositide and cholesterol

**DOI:** 10.64898/2025.12.29.696921

**Authors:** Jie Li, Jay Xiaojun Tan, Zhijian J. Chen, Xuewu Zhang, Xiao-chen Bai

## Abstract

Stimulator of interferon genes (STING) is an essential adaptor in the cytosolic DNA sensing innate immune pathway. STING is activated by cyclic-GMP-AMP (cGAMP) produced by the DNA sensor cGAMP synthase (cGAS). cGAMP-induced high-order oligomerization and translocation of STING from the endoplasmic reticulum to Golgi and post-Golgi vesicles are critical for STING activation. Recent studies have shown that phosphatidylinositol phosphates (PIPs) and cholesterol also play important roles in STING activation, but the underlying mechanisms remain unclear. Here, we demonstrate that cGAMP-induced high-order oligomerization of STING is enhanced strongly by PI(3,5)P_2_ and PI(4,5)P_2_, and by PI(4)P to a less extent. Our cryo-EM structures reveal that PIPs together with cholesterol bind at the interface between STING dimers, directly promoting the high-order oligomerization. The structures also provide an explanation for the preference of the STING oligomer to different PIPs. Mutational and biochemical analyses confirm the binding modes of PIPs and cholesterol and their roles in STING activation. Our findings shed light on the regulatory mechanisms of STING by specific lipids, which may underlie the role of intracellular compartment trafficking in dictating STING signaling.

## Main

The cGAS/STING pathway serves as a fundamental mechanism to detect cytosolic foreign DNA from invading pathogens, initiating innate immune responses against infections^1^. Activated by DNA, cGAS (cGAMP synthase) generates the second messenger 2’,3’-cyclic-GMP-AMP (cGAMP), which binds to and activates the adaptor protein STING (Stimulator of Interferon Genes)^2–5^. STING initiates multiple downstream pathways, including interferons, NF-κB, autophagy, and inflammasome, to eliminate pathogens. In addition, the cGAS/STING signaling pathway plays a pivotal role in anti-tumor immunity, due to activation of the pathway by fragmented DNA as a result of chromosomal instability in cancer cells and detection of tumor DNA in dendritic cells^1^. Recent studies have shown that, depending on the context, STING-mediated signaling can either positively or negatively regulate tumor development and metastasis^6–9^. On the other hand, abnormal activation of STING could cause autoimmune diseases, age-related inflammation and neurodegeneration^10,11^. Various agonists and antagonists of STING are actively investigated as potential therapeutics for viral infection, cancer and neurodegenerative diseases.

STING is an endoplasmic reticulum (ER) membrane protein comprising four transmembrane (TM) helices forming the TM domain (TMD), a cytoplasmic ligand-binding domain (LBD) responsible for cGAMP binding, and a C-terminal tail containing both the phosphorylation site (Ser366) and the PXPLRXD (single-letter amino acid code; X, any amino acid) motif for recruiting TANK-binding kinase 1 (TBK1)^12–15^. STING forms a constitutive domain-swapped dimer, which adopts an inactive conformation in the absence of cGAMP. Binding of cGAMP to the LBD induces a set of concerted conformational changes in STING, leading to its high-order oligomerization, which is essential for the activation of the pathway^12,13,16,17^. TBK1 binding to the C-terminal tails in the STING oligomers facilitates the phosphorylation of Ser366^12,14^. Phosphorylated STING recruits and promotes TBK1-mediated phosphorylation of the transcription factor interferon regulatory factor 3 (IRF3), triggering type I interferon expression. Previous structural studies have elucidated the mechanisms by which the high-order oligomerization of STING serves as a platform, facilitating efficient recruitment of TBK1 as well as the phosphorylation of both STING and IRF3 by TBK1^12–14,16–18^.

In addition to the high-order oligomerization, STING activation requires its translocation from ER to the Golgi apparatus and downstream compartments^19,20^. Elevated ER exit of STING can trigger its spontaneous activation in the absence of cGAMP, leading to immune dysregulation that underlies diseases such as COPA syndrome^21,22^. Lipids that selectively distribute to distinct membrane compartments may regulate the activation of STING, which could provide one possible explanation for the crucial role of the ER-to-Golgi translocation in STING activation. Indeed, several lines of evidence suggest that STING may directly interact with and be modulated by phosphatidylinositol phosphates (PIPs) and cholesterol, which exhibit different distributions in the ER, Golgi and endosomes^23–30^. Nonetheless, the precise mechanisms by which these lipids interactions influence STING activity remain unclear.

Here, we show that both PI(3,5)P_2_ and PI(4,5)P_2_ can substantially enhance the high-order oligomerization of cGAMP bound STING. Our high-resolution cryo-EM structures reveal that PIPs bind the interfaces between two STING dimers, acting effectively as bridging molecules that promote the high-order oligomerization of STING. Interestingly, the structures show that this interaction is stabilized by a cholesterol-like molecule, positioned between STING and PIPs. Our mutational analyses, lipid binding, in vitro liposome reconstitution, biochemical and cell-based experiments confirmed these binding modes of PIPs and cholesterol, and their roles in promoting high-order STING oligomerization and activation. These results, together with our co-submitted manuscript (Tan et al), reveal an additional layer of the regulatory mechanism governing STING activity that potentially underlies the requirement of the ER-Golgi translocation in STING signaling.

### PIPs enhance cGAMP-induced high-order oligomerization of STING

To investigate the role of PIPs in STING high-order oligomerization, we first examined whether commonly found PIPs, such as PI(4)P, PI(3,5)P_2_, and PI(4,5)P_2_, could induce STING oligomerization through a native gel assay. Consistent with our published findings^16,18^, STING primarily existed as a dimer in the absence of any ligand (Figure 1a). When treated with cGAMP, PI(3,5)P_2_, or PI(4,5)P_2_ individually, STING formed tetramers and higher-order oligomers. Dose-response analysis indicated that PI(3,5)P_2_ has a slightly stronger effect than PI(4,5)P_2_ in promoting oligomerization (Extended Data Fig. 1a). In contrast, PI(4)P was much less effective in inducing STING oligomerization (Figure 1a). When combined with cGAMP, PI(3,5)P_2_ or PI(4,5)P_2_ enhanced the formation of larger STING oligomers (Figure 1a). Co-treatment with the recently discovered STING agonists STG2 or C53^16,18,31^, which binds two distinct sites in the TMD of STING^16,18,31^, further increased the high-order oligomerization of STING (Figure 1a and b). Consistently, cryo-EM images showed that STING bound to either cGAMP/PI(3,5)P_2_/STG2 or cGAMP/PI(4,5)P_2_/STG2 formed abundant long oligomers, through side-by-side packing of multiple dimers with an characteristic overall positive curvature (Extended Data Fig. 1b), similar to those of STING bound to cGAMP/C53/STG2 seen previously^18^. In contrast, cGAMP/PI(4)P/STG2 induced much less and shorter oligomers of STING (Extended Data Fig. 1b). These findings together indicate that PI(3,5)P_2_ and PI(4,5)P_2_, but not PI4P, are effective enhancers of cGAMP-induced high-order oligomerization of STING.

**Figure 1.**
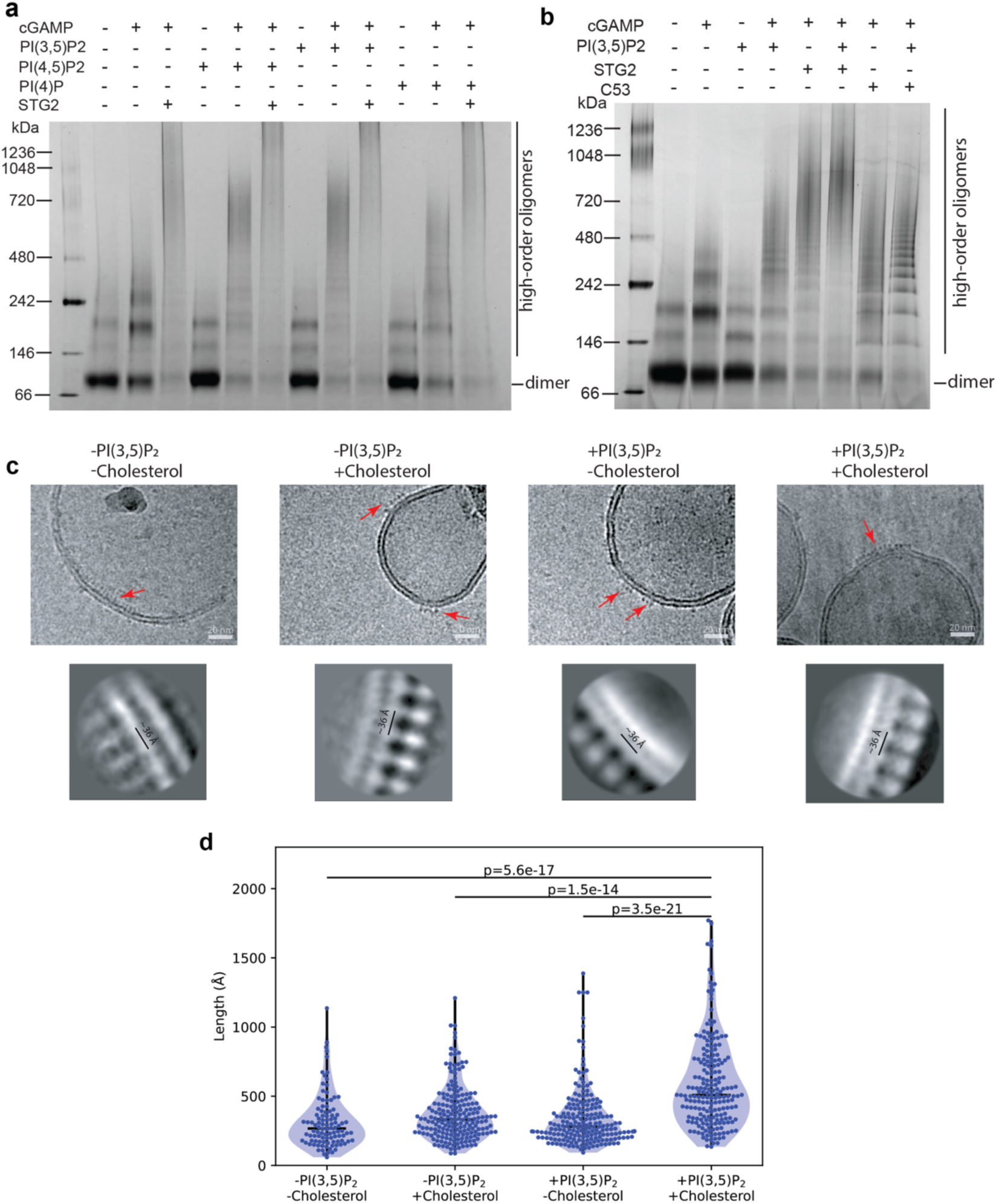
PIPs enhance the high-order oligomerization of STING. (**a**) and (**b**) Analyses of the high-order oligomerization of STING treated with different combinations of agonists and PIPs by native PAGE. Images shown are representatives from three biological repeats. (**c**) High-order oligomers of cGAMP-bound STING on liposomes containing different combinations of PI(3,5)P_2_ and cholesterol. The upper and lower panels show motion-corrected micrographs and 2D class averages of STING oligomers, respectively. (**d**) Length distribution of STING oligomers on liposomes with different combinations of PI(3,5)P_2_ and cholesterol. The numbers of micrographs collected for the -PI(3,5)P_2_/-cholesterol, -PI(3,5)P_2_/+cholesterol, +PI(3,5)P_2_/-cholesterol, +PI(3,5)P2/+cholesterol conditions were 1,204, 5,211, 1,971, and 2,862, respectively. Each dot represents one filament. The horizontal bars represent the medians. P-values were calculated using the two-sided Mann-Whitney U test.

### PIPs and cholesterol together promote the oligomerization and phosphorylation of STING in lipid membrane

We next reconstituted STING into liposomes to examine the roles of PI(3,5)P_2_ and cholesterol in its oligomerization in lipid membrane. Cryo-EM micrographs showed that cGAMP-bound STING formed high-order oligomers on the surface of liposomes (Figure 1c). The length of STING filaments on liposomes containing both PI(3,5)P_2_ and cholesterol are significantly longer than those on liposomes missing either one or both lipids (Figure 1d). In addition, 2D class averages of the oligomers from liposomes with both PI(3,5)P_2_ and cholesterol are shaper than the others (Figure 1c). These results suggest that PI(3,5)P_2_ and cholesterol together strongly enhanced the high-order oligomerization of STING. Interestingly, while most STING oligomers on liposomes containing PI(3,5)P_2_ and cholesterol or either one are located on the outer surface of liposomes and adopt positive curvature similar to those in detergent solution, oligomers on liposomes without these two lipids often sit on the inner surface of liposome and show negative curvature (Figure 1c). These observations suggest that PI(3,5)P_2_ and cholesterol strengthen the interactions between the STING TMD and thereby tighten the TMD portion of the oligomer.

Using liposome-reconstituted STING, we further investigated the regulation of TBK1-mediated phosphorylation of STING by PIPs and cholesterol (Figure 2a). PI(3,5)P_2_, PI(4,5)P_2_ but not PI4P enhanced cGAMP-stimulated phosphorylation of STING by TBK1 (Figure 2b). We then focused on PI(3,5)P_2_ and cholesterol to check their effects on regulation STING phosphorylation. The results showed that the phosphorylation of STING upon cGAMP stimulation was substantially enhanced by the presence of 10% cholesterol or 5% PI(3,5)P_2_ in liposomes (Figure 2c). The phosphorylation levels were further increased when both lipids were present (Figure 2d). The optimal concentration of cholesterol was ∼10%, while cholesterol at either 5% or 20% led to lower levels of STING phosphorylation (Figure 2e). PI(3,5)P_2_ also enhanced the phosphorylation level at the condition with 20% cholesterol, further demonstrating the synergistic effect between cholesterol and PI(3,5)P_2_ on STING activation (Figure 2f).

**Figure 2.**
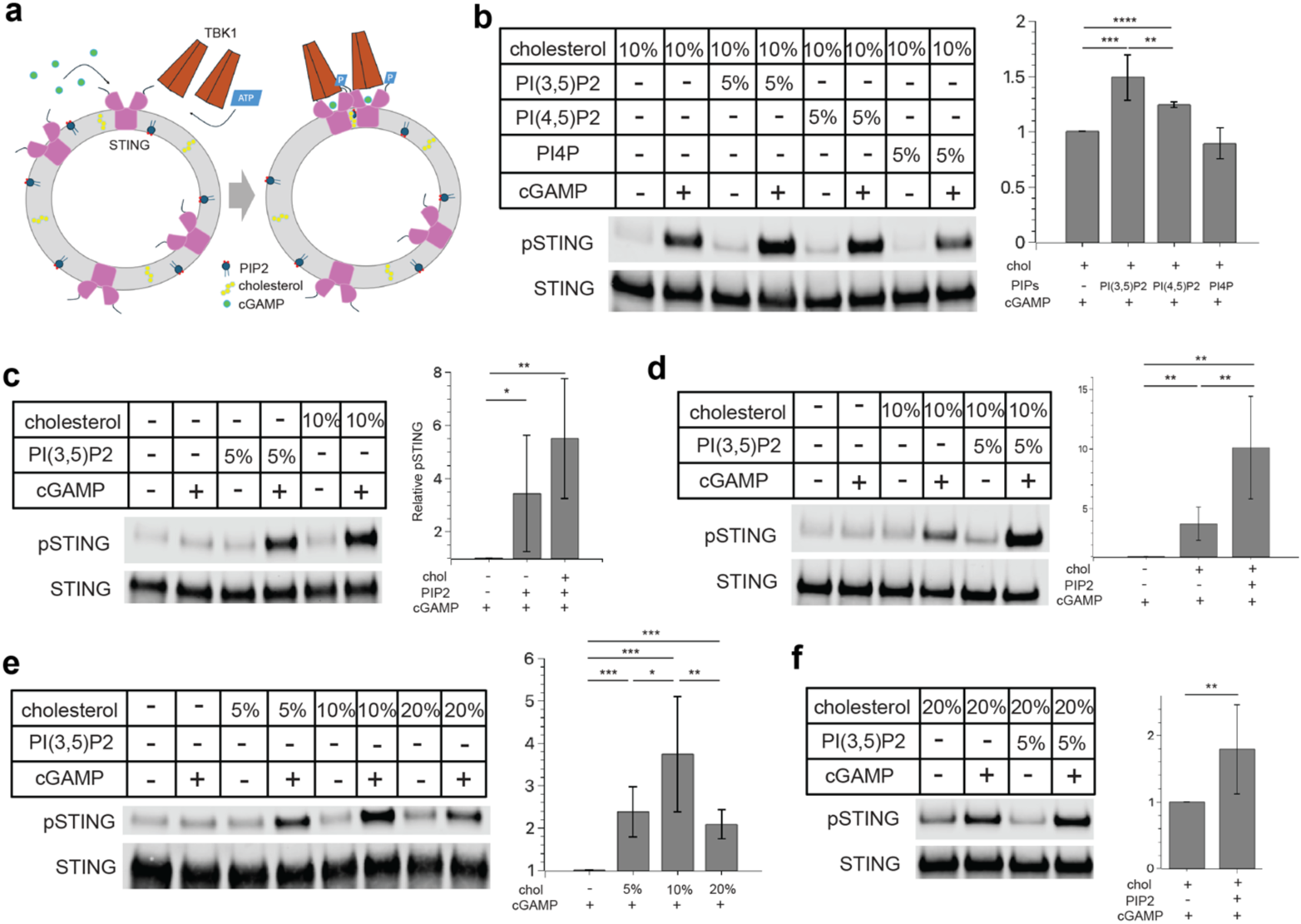
Effect of PIPs and cholesterol on STING phosphorylation by TBK1 on liposomes. (**a**) Schematic diagram of cGAMP-induced STING phosphorylation by TBK1 on liposomes. (**b**) Comparison of the effects of PI(3,5)P_2_, PI(4,5)P_2_ and PI4P on STING phosphorylation by TBK1 on liposomes containing 10% cholesterol. (**c**) Comparison of the effects of cholesterol and PI(3,5)P_2_ on STING phosphorylation by TBK1 on liposome. pSTING level is increased more than 2-fold by cholesterol or PI(3,5)P_2_. (**d**) cGAMP, PI(3,5)P_2_ and cholesterol together induce higher levels of STING phosphorylation by TBK1 on the liposome than does each ligand individually. pSTING level is increased more than 5-fold by cholesterol and PI(3,5)P_2_ together. (**e**) Comparison of STING phosphorylation by TBK1 on liposome with various concentration of cholesterol. pSTING level is increased around 2-fold by 5% or 20% cholesterol while increased around 4-fold by 10% cholesterol. (**f**) Effect of PI(3,5)P_2_ on STING phosphorylation by TBK1 on liposomes containing 20% cholesterol. The observations in (**b-f**) are mean ± s.d from three or four biological repeats. P-values in (**b-f**) were calculated with two-sided paired Student’s t-test.

### Cryo-EM structures of STING bound to PI(3,5)P_2_ or PI(4,5)P_2_ at high resolution

To understand how PI(3,5)P_2_ and PI(4,5)P_2_ promote the high-order oligomerization of STING, we sought to determine the cryo-EM structures of STING in the presence of these PIPs (Figure 3a-c). To facilitate the cryo-EM structural determination, both cGAMP and STG2 were added into the sample to increase the structural stability of the STING oligomer. We divided the long oligomers into small segments containing 6 dimers of STING for 3D reconstruction. This approach allowed us to determine the cryo-EM structures of the STING/PI(3,5)P_2_ and STING/PI(4,5)P_2_ oligomers at a resolution of 2.8 Å and 2.6 Å, respectively (Extended Data Fig. 2 and 3). We built atomic models of the tetrameric unit to the center portion of the maps, which could be extended according to the inter-dimer packing mode to reconstruct higher-order oligomers. The structures unveiled that each STING dimer assumes the conformation similar to STING bound with cGAMP, C53 and STG2^18^, suggesting that PIPs binding does not induce any conformational change within the STING dimer (Figure 3d and e).

**Figure 3.**
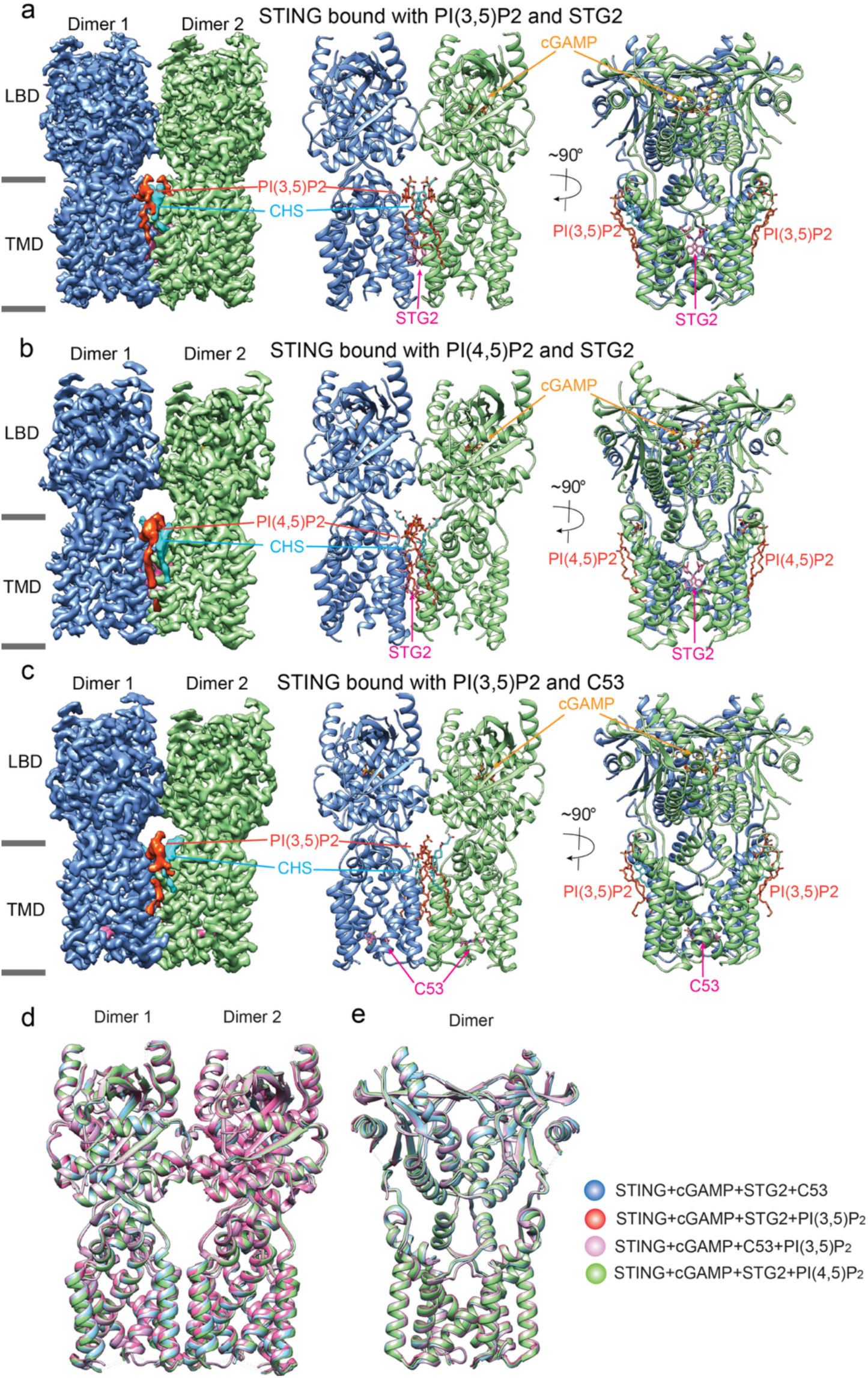
Overall structures of the STING high-order oligomer bound to different PIPs and agonists. (**a**) Cryo-EM map and atomic model of the STING high-order oligomer bound to cGAMP, PI(3,5)P_2_ and STG2. (**b**) Cryo-EM map and atomic model of the STING high-order oligomer bound to cGAMP, PI(4,5)P_2_ and STG2. (**c**) Cryo-EM map and atomic model of the STING high-order oligomer bound to cGAMP, PI(3,5)P_2_ and C53. (**d**) and (**e**) Comparison of the structures of the high-order oligomer of STING bound to different combinations of PIPs and agonists.

In both the cryo-EM maps of STING/PI(3,5)P_2_ and STING/PI(4,5)P_2_ tetramers, robust additional densities appear on the cytosolic side of STING-TMD between two dimers. These densities could be attributed to PI(3,5)P_2_ and PI(4,5)P_2_, respectively, based on clear features consistent with the head groups (Figure 3a and b, Figure 4). While one of two acyl chains of PI(3,5)P_2_ and PI(4,5)P_2_ could be modeled into the cryo-EM density, the second acyl chain of PIPs was poorly resolved in the cryo-EM map due to flexibility and thus could not be accurately modeled. Moreover, an extra density proximal to PIPs was observed in both cryo-EM maps, exhibiting a flat and elongated shape reminiscent of cholesterol. Given the presence of cholesteryl hemisuccinate (CHS) in the buffer, we attributed this additional density to CHS (Figure 3a and b, Figure 4). This observation supports our in vitro liposome reconstitution results showing that cholesterol works cooperatively with PIPs to promote the high-order oligomerization of STING. Notably, the packing interaction between the two STING dimers in these structures in the presence of PIPs is nearly identical to that observed in the previous cGAMP/STG2/C53-induced STING oligomer in the absence of PIPs^16,18^ (Figure 3d and e). This structural comparison supports the idea that this mode of STING oligomerization represents the native high-order oligomeric state of STING in the cell.

**Figure 4.**
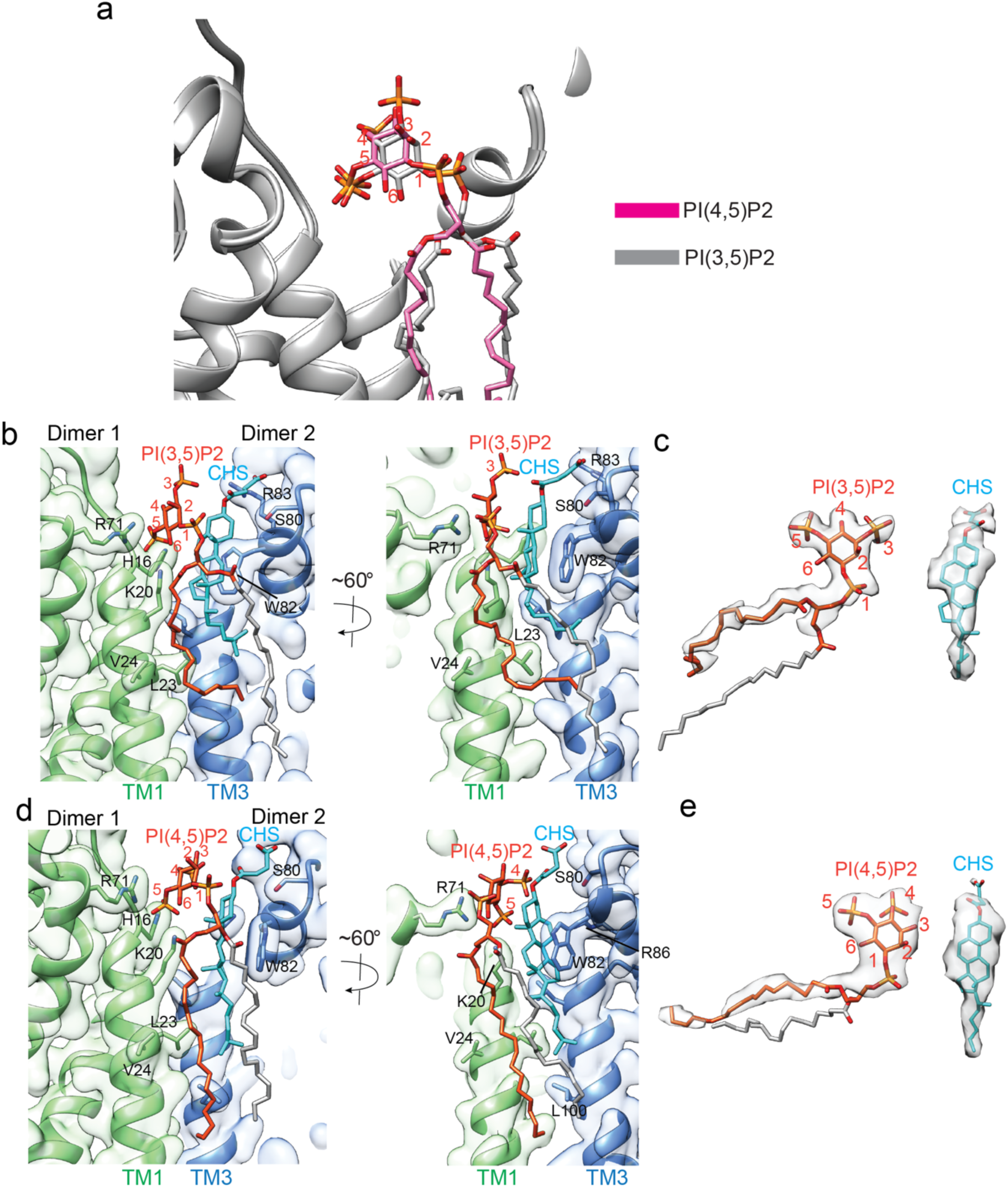
PIPs and CHS bind at the interface between two STING dimers. (**a**) Comparison of the binding modes of PI(3,5)P_2_ and PI(4,5)P_2_. (**b**) Detailed view of the binding modes of PI(3,5)P_2_ and CHS at the interface between two STING dimers in the structure of STING bound to cGAMP/PI(3,5)P_2_/STG2. (**c**) Cryo-EM density of PI(3,5)P_2_ and CHS in the structure of STING bound to cGAMP/PI(3,5)P_2_/STG2. (**d**) Detailed view of the binding modes of PI(4,5)P_2_ and CHS at the interface between two STING dimers in the structure of STING bound to cGAMP/PI(4,5)P_2_/STG2. (**e**) Cryo-EM density of PI(4,5)P_2_ and CHS in the structure of STING bound to cGAMP/PI(4,5)P_2_/STG2. In **b**-**e**, the acyl chain of PIPs without clear density is shown in gray.

Furthermore, we determined the cryo-EM structure of STING in the presence of cGAMP/PI(4)P/STG2 (Extended Data Fig. 4). While the overall resolution of this structure (2.8 Å) is similar to those bound to PI(3,5)P_2_ or PI(4,5)P_2_, PI(4)P and CHS show rather weak density and therefore not included in the atom model (Extended Data Fig. 5a and b). This observation is consistent with the native gel results showing that PI4P is not as strong as PI(3,5)P_2_ and PI(4,5)P_2_ in promoting the high-order oligomerization of STING.

To exclude the possibility that the observed binding mode of PIPs might be an artifact as a result of the STG2 binding, we also determined the cryo-EM structure of STING/cGAMP/PI(3,5)P_2_ in complex with C53 (Extended Data Fig. 6). The cryo-EM structures revealed that the binding modes of PI(3,5)P_2_ in both STG2-bound and C53-bound STING are virtually identical (Figure 3c and Extended Data Fig. 6e and f). Finally, we determined the cryo-EM structure of STING in the presence of only cGAMP and PI(3,5)P₂, without either STG2 or C53. The cryo-EM micrographs of this sample revealed similar but shorter STING oligomers. The cryo-EM map at 4.1 Å resolution shows that PI(3,5)P₂ is bound to the same site on STING (Extended Data Fig. 7). These results together strongly support the notion that this mode of PIP binding is an inherent regulatory mechanism of STING, not an artifact induced by the TMD agonists.

### Detailed binding mode of PI(3,5)P_2_ and PI(4,5)P_2_

In the structures of the STING/PI(3,5)P_2_ and STING/PI(4,5)P_2_ complexes, the PIP molecule is located within the same groove between the TMDs of two adjacent STING dimers (Figure 4a). Specifically, the 5-phosphate group of either PI(3,5)P_2_ or PI(4,5)P_2_ interacts with residues H16 and K20 of TM1 and R71 of TM2 within one STING dimer (Figure 4b-e). The inositol ring and a part of the acyl chains of PI(3,5)P_2_ or PI(4,5)P_2_ pack against one side of the CHS molecule, which in turn uses its other side to make stacking interactions with W82 from the neighboring STING dimer (Figure 4b and d). One of the acyl chains of PI(3,5)P_2_ and PI(4,5)P_2_ also contacts several hydrophobic residues in STING, including L23 and V24 in TM1 (Figure 4b and d). Moreover, S80 in the second STING molecule is placed in close proximity to the hydrophilic region of CHS (Figure 4b and d). This binding configuration of PI(3,5)P_2_/CHS and PI(4,5)P_2_/CHS enhances the interaction between the two STING dimers, thereby promoting the formation of higher-order STING oligomers.

It is worth pointing out that 2-OH group in the inositol ring of PIPs is positioned axially, perpendicular to the ring plane. In our structures, the inositol ring is orientated so that the 2-OH points outwards, while the opposite side of the ring packs against CHS (Figure 4). This arrangement positions the 5-phosphate group to interact with the positively charged residues in STING. A 180°-flip of the inositol ring of PI(3,5)P_2_ along the C1-C4 axis could, in principle, place the 3-phosphate in this position to engage STING. However, this would orient the 2-OH towards CHS and potentially cause steric clashes between PI(3,5)P_2_ and CHS. These analyses suggest a structural basis for the requirement of the 5-phoshate group in PIPs for binding STING. On the other hand, the 4-phosphate group is located further away and unlikely contribute much to the interaction with STING. This, along with its reduced charge compared with PI(3,5)P_2_ and PI(4,5)P_2_, explains why PI4P is a weaker binder of STING.

### Validation of the PIP binding mode using lipid binding and native PAGE assays

To validate the PIP binding mode observed in the cryo-EM structures, we mutated residues in STING surrounding PIP and CHS, including K20, R71, S80 and W82, and tested their effects on the direct binding to PIPs and cholesterol using a lipid-strip assay^32^. Our results show that STING wild type (WT) binds to most PIP lipids, cholesterol and its derivatives, but not several other types of lipids such as non-phosphorylated phosphatidylinositol (PI), phosphatidylethanolamine (PE) and phosphatidylserine (PS). The R71H/K20E mutation abolishes binding to PIPs, while the S80I mutant exhibits reduced binding to cholesterol compared with STING WT (Extended Data Fig. 8). These results support the binding mode of PIPs and cholesterol shown in our structures, although we could not evaluate the preference of STING for different PIPs due to the non-quantitative nature of this assay.

We next examined high-order oligomerization of these STING mutants in response to PIPs using the native page assay. Purified STING S80I and W82Q mutants exhibit greatly reduced ability of forming large oligomers in response to either PI(3,5)P_2_ or PI(4,5)P_2_ (Figure 5a). STING K20E still robustly formed large oligomer when stimulated by PI(3,5)P_2_ or PI(4,5)P_2_ (Figure 5a), which is consistent with the fact that the interaction between lysine and phosphate is in general weaker than that between arginine and phosphate^33^. The STING R71H mutant exhibits a moderately reduced ability to form large oligomers, while the K20E/R71H double mutation led to a much more pronounced reduction in STING oligomer formation (Figure 5a). Furthermore, we performed native PAGE on cell lysates expressing STING WT and the K20E/R71H and S80I mutants. Both mutants showed markedly reduced ability to form higher-order oligomers upon cGAMP stimulation in vivo, validating the binding mode between PIPs and STING as seen in the cryo-EM structures (Figure 5b). Notably, the S80I mutant but not WT STING appeared as a smeared band on native PAGE, whereas purified S80I displayed a sharp peak on size-exclusion chromatography (SEC) and eluted at a position similar to the WT (Extended Data Fig. 9a). These observations suggest that the S80I mutant is properly folded but has reduced structural stability compared to the WT protein. Purified protein S80I and W82Q also showed an additional band between dimer and tetramer on native PAGE, possibly arising from protein instability (Figure 5a). Furthermore, density corresponding to CHS was observed in our previously resolved STING oligomer structures in the absence of PIPs (Extended Data Fig. 9b). Together, these findings support an idea that cholesterol not only promotes higher-order oligomerization but also acts as a structural component that contributes to STING structural stability.

**Figure 5.**
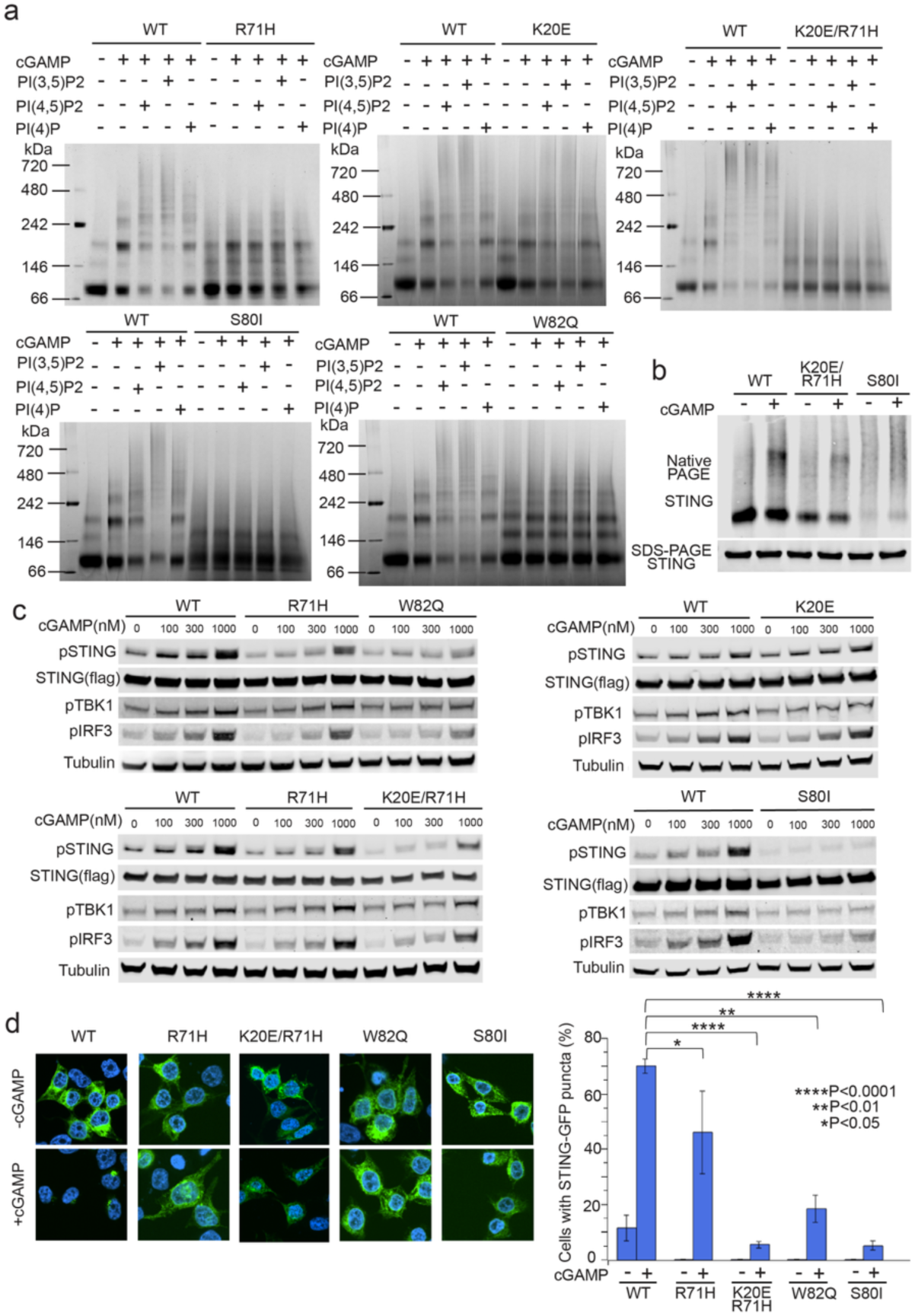
Mutational analyses of the binding interface in STING for PIPs and cholesterol. (**a**) Mutations in the binding interface compromise the high-order oligomerization of purified STING in the native PAGE assay. The images shown are representatives of three biological repeats. (**b**) Mutations in the binding interface compromise the high-order oligomerization of STING in HEK293T cells. Same samples were loaded to native PAGE and SDS-PAGE separately. Both the gels were blotted using the same STING antibody. (**c**) Western blots showing that mutations in the binding interface reduce cGAMP-stimulated phosphorylation of STING, TBK1 and IRF3 in HEK293T cells. (**d**) Mutations in the binding interface reduce cGAMP-indued puncta formation of STING-GFP in HEK293T cells. The images shown are representatives of three biological repeats. Scale bars, 5 μm. Cells transfected with each STING construct were imaged in a blind manner in three independent experiments. The percentage of cells with STING-GFP puncta was calculated based on 150∼250 GFP-positive cells for each construct in each experiment. Specifically, 206, 279, 114, 255, and 183 cells for WT, R71H, R71H/K20E, W82Q, S80I in the 1^st^ experiment; 203, 126, 139, 216, and 194 cells in the 2^nd^ experiment; 180, 171, 115, 207, and 178 cells in the 3^rd^ experiment. Statistics data are mean ± s.d. P-values were calculated with two-tailed Student’s t-test at a 95% confidence level.

### Functional evaluation of the role of PIPs in STING activation using cell-based assay

The results of the lipid binding, native gel assay and cryo-EM structures support the model that PIPs and cholesterol directly bind STING and enhance its activation upon stimulation by cGAMP in cells. To test this model, we used cell-based experiments to evaluate how mutations that disrupt the binding of PIPs and cholesterol affect STING signaling. Our cell-based functional assay revealed that mutations to residues involved in the interactions with PIPs and cholesterol, including R71H, K20E/R71H, S80I and W82Q, substantially reduced phosphorylation of STING, TBK1 and IRF3 triggered by cGAMP (Figure 5c). The strong reduction of phosphorylation as a result of mutations of S80 and W82, two residues that are important for CHS binding, further supports the involvement of cholesterol in the cGAMP-mediated activation of STING (Figure 5c).

Besides phosphorylation, another hallmark of cGAMP-induced activation of STING in cells is that it translocates from the ER to the Golgi apparatus and post-Golgi vesicles, where it forms large puncta as a result of the high-order oligomerization. Our fluorescent images showed that the K20E/R71H, S80I and W82Q mutants of STING expressed at similar levels to WT STING prior to cGAMP treatment (Figure 5d). cGAMP treatment led to robust puncta formation in cells expressing WT STING, whereas no large puncta were observed in cells expressing the mutants (Figure 5d). These findings together support our structure-based model in which PIPs and cholesterol directly bind STING and contribute to its high-order oligomerization and activation in cells.

## Discussion

The findings in this study suggest that phosphatidylinositol phosphate lipids and cholesterol can directly bind to STING, and thereby enhancing its high-order oligomerization and activation. Our results showed that both PI(3,5)P_2_ and PI(4,5)P_2_ bind the same site on STING and have similar potency in inducing the high-order oligomerization, but PI(4)P is much less potent. The lower level of sensitivity of STING towards PI(4)P might be a safeguard mechanism to avoid spontaneous activation of STING in the absence of upstream signal, considering that PI(4)P is highly abundant in the Golgi membrane^34^. A recent study has shown that STING activation can be altered by manipulating the PI(4)P levels in Golgi through either pharmacological or genetic approaches, suggesting that PI(4)P plays a role in STING regulation^29^. It is possible that this effect of PI(4)P is indirect, as changes in the concentration of PI(4)P could affect other PIP species such as PI(4,5)P_2_ and PI(3,5)P_2_. Oxysterol-binding protein (OSBP) can exchange PI(4)P and cholesterol between the ER and Golgi membranes, suggesting that PI(4)P may also play a role in regulating cellular cholesterol distribution, which thereby indirectly regulates the STING signaling. PI(4,5)P_2_ is enriched in the plasma membrane, but also present in the Golgi and endosomes at much lower levels^34,35^. PI(3,5)P_2_ is mostly localized to late endosomes and lysosomes^34^. Upon cGAMP binding, STING traffics to Golgi and Golgi-derived endosomes, but not plasma membrane. In the co-submitted manuscript (Tan et al), we found that STING constitutively interacted with PIKfyve, a kinase that phosphorylates PI3P to produce PI(3,5)P_2_, and PI(3,5)P_2_-binding mutants of STING showed defects in ER-to-Golgi trafficking. These results suggest that low levels of PI(3,5)P_2_ in the ER generated by STING-bound PIKfyve is required for the ER-exit of STING.

Our cryo-EM structures suggest that PIPs and cholesterol in an inter-dependent manner strengthen the dimer-dimer interaction in the STING oligomer, supporting previous studies that cholesterol plays a role in STING activation and regulation^23–25,30^. Interestingly, re-examination of our earlier cryo-EM structure of the STING oligomer in the absence of PIPs revealed a cholesterol-like density at the same site identified in the present work (Extended Data Fig. 9b), suggesting that cholesterol binding could occur independently of PIP binding. Because both the ER and Golgi contain certain levels of cholesterol, it is plausible that STING binds cholesterol in both organelles. Cholesterol may play distinct roles at different stages of STING regulation. At the ER, cholesterol binding, together with other factors such as STIM1, may help retain STING within the ER^36,37^. In contrast, our data suggest that cholesterol acts synergistically with PIPs to promote STING oligomerization and activation at Golgi or post-Golgi compartments.

Our lipid binding, native PAGE and cryo-EM studies together indicate that both PI(3,5)P_2_ and PI(4,5)P_2_ are able to strongly bind STING and effectively induce high-order STING oligomerization. Considering the distribution patterns of PI(3,5)P_2_ and PI(4,5)P_2_, the sites of STING activation and signaling could include post-Golgi endolysosomes, in addition to the ER-Golgi intermediate compartments and the Golgi apparatus as previously suggested^1,19,20^. While both PI(3,5)P_2_ and PI(4,5)P_2_ show similar binding modes and potencies in inducing high-order oligomers of STING, PI(3,5)P_2_ may play a more important role in STING activation in cells considering its higher abundance in late endosomes. In addition, PIKfyve bound to STING may finely tune the amounts of PI(3,5)P_2_ on local membranes to help precisely control STING trafficking and signaling in response to infection (Tan et al, co-submitted).

R71 of STING forms a salt bridge with the 5-phosphate group of PI(3,5)P_2_ or PI(4,5)P_2_. Our mutational analyses confirmed that R71 is crucial for the binding and activating effect of PIPs on STING. It is worth mentioning that R71H is a part of the STING HAQ (R71H-G230A-R293Q) variant, which is one of the prevalent human STING alleles^38,39^. The HAQ variant has impaired immune functions, which has been largely attributed to the R71H mutation^38,39^. Our cryo-EM results provide a structural explanation for the critical role of R71 in STING activation, which likely underlies the detrimental effect of the R71H mutation on STING-mediated immunity.

In summary, our findings reveal the mechanistic basis for the pivotal role of PIPs and cholesterol in the activation of STING, opening avenues for understanding the molecular details that underlie the role of intracellular compartment trafficking in dictating STING function.

## Materials and methods

### Lipid stock preparation

Lipids including 18:1 PI(3,5)P_2_, 18:1 PI(4,5)P_2_, 18:1 PI(4)P were purchased from Avanti Polar lipids. To make lipid solutions, desired amounts of lipid chloroform stock were dispensed to a glass culture tube and dried by nitrogen gas stream. lipids were further dried by high vacuum overnight and solubilized in a detergent buffer containing 20 mM HEPES-Na, pH 7.4, 150 mM NaCl, 0.03% DDM, 0.003% CHS to a final concentration of 1 mM.

### Protein expression and purification

Similar to previous studies, the coding sequence of human STING (1-343) followed by an HRV-3C protease recognition sequence and the purification tag T6SS immunity protein 3 (Tsi3) coding sequence was inserted into the pET-BM vector^13,18^. The protein was expressed in Freestyle 293-F cells either by baculovirus infection or transfection using polyethylenimine (PEI). Mutations were introduced by Q5 mutagenesis kit (New England Biolabs). Cells were collected 50 hrs after transfection and resuspended in buffer A containing 20 mM Tris-HCl, pH 8.0, 200 mM NaCl, 1 mM DTT and protease inhibitor cocktail (Roche). After disrupting the cells by high-pressure cell disruptor, the membrane fraction was spun down by ultracentrifugation at 150,000g for 1 hr. Membrane proteins were extracted by buffer A supplemented with 1% (w/v) n-dodecyl-β-D-maltopyranoside (DDM) and 0.2% cholesteryl hemisuccinate Tris salt (CHS) (Anatrace). After ultracentrifugation at 100,000 g for another 1 hr, the supernatant was supplemented with 1 mM CaCl_2_. The STING protein was captured by Tse3-conjugated Sepharose resin. The wash buffer contained buffer A supplemented with 0.06% DDM, 0.006% CHS and 1 mM CaCl_2_. STING was eluted by HRV-3C cleavage and further purified by size-exclusion chromatography in buffer B (20 mM HEPES-Na, pH 7.4, 150 mM NaCl, 0.03% DDM and 0.003% CHS, 1 mM DTT). The main peak was pooled and incubated with cGAMP (Cayman chemical), PIP lipids and STG2 or C53 at 1:2:2:2 molar ratio for 3 hrs. The samples were concentrated to 4-6 mg/ml for cryo-EM grid preparation.

### Native gel assay

Purified STING wild type or mutants at 20 µM in buffer B was incubated with different combinations of the compounds cGAMP, PIP lipids, C53 and NVS-STG2 at 40 µM at 4 °C for 3 hrs. The samples were mixed with the native gel sample buffer, 0.25% G-250 sample additive and resolved by a 3–12% gradient native gel (Invitrogen, cat. #BN1003BOX).

### Protein-lipid interaction assay by lipid strips

Membrane lipid strips (Cat. # P-6002) and PIP arrays (Cat. # P-6100) were purchased from Echelon Biosciences. To prepare cholesterol strips, 1 µL aliquots of cholesterol dissolved in chloroform at defined concentrations were applied onto PVDF membranes. Blotting was performed according to the manufacturer’s protocol. 3 ug/ml STING(1-343) protein was incubated with strips or arrays. The binding was detected with antibody against STING.

### Cryo-EM data collection and image processing

The samples of STING in complex with cGAMP/STG2/PI(3,5)P_2_, cGAMP/STG2/PI(4,5)P_2_, cGAMP/STG2/PI(4)P, or cGAMP/C53/PI(3,5)P_2_ were applied to glow-discharged Quantifoil R1.2/1.3 300-mesh gold holey carbon grids (Quantifoil, Micro Tools GmbH, Germany). Grids were blotted under 100% humidity at 4 °C and plung-frozen in liquid ethane using a Mark IV Vitrobot (FEI). Micrographs were collected in the super-resolution mode on Titan Krios microscopes (FEI) with K3 Summit direct electron detectors (Gatan). The nominal magnification and pixel size of each data set are summarized in Tables S1.

Motion-correction and dose-weighting of the micrographs were carried out using the Motioncor2 program (version 1.2)^40^. GCTF 1.06 was used for CTF correction^41^. Template-based particle picking of segments of the STING high-order oligomers was carried out using the autopick tool in the single particle mode in RELION 4.0^42^. Particles were extracted with box sizes large enough to contain 6 dimers in the STING oligomers. To maximize the number of particles, particles were also picked with crYOLO (version 1.9) for three of the four datasets ^43^. Particles from both picking methods were cleaned up with multiple rounds of 2D and 3D classification in RELION. Good particles from the two methods were joined with duplicated removed and subjected to 3D refinement with C2 symmetry. During 2D and 3D classification steps, shorter oligomers were removed, leading to reconstructions of at least four dimers. The exact procedures were different to some extent for each dataset, which are summarized in Extended Data Fig. 2, 3, 4 and 6. The cryo-EM map of the STING tetramer in complex with cGAMP/C53 (EMD ID: EMD25142) was low-pass filtered and used as the initial mode for 3D classification and refinement. The refined maps were further improved by using Bayesian polishing and CTF refinement at the final stage. The Fourier Shell Correlation (FSC) 0.143 criterion was used for estimating the resolution of the maps. Local resolution was calculated in RELION.

### Model building and refinement

To build the atomic models of the structures with different ligands bound, the published model of human STING tetramer bound to cGAMP, STG2 and C53 (PDB ID: 8FLM) was docked into the cryo-EM maps. The models were adjusted manually, and the corresponding PIP ligands and agonists were added in Coot 0.98^44^. PI(4)P or CHS was not included in the model for the STING/ cGAMP/STG2/PI(4)P) map due to lack of clear density, while PIP lipids and CHS were clearly resolved and therefore modeled in the other three structures. The models were refined using the real-space refinement module in Phenix 1.18 and ISOLDE 1.7 as a plugin tool in ChimeraX 1.7 ^45–47^. Model quality was checked using Molprobity as a part of the Phenix validation tool set^48^. Model statistics are summarized in Table S1. Structural figures were rendered in Chimera 1.16 or ChimeraX 1.7^46,49^.

### Antibodies

The following antibodies were used: rabbit anti-STING antibody (Cat. #13647S), rabbit anti-phospho-STING (S366) antibody (Cat. #19781S), rabbit anti-phospho-TBK1/NAK(S172) antibody (Cat. #5483S), mouse anti-flag antibody (Cat. #8146S) and mouse anti-α-Tubulin antibody (Cat. #3873S) were purchased from Cell signaling Technology; rabbit anti-phospho-IRF3(S386) (Cat. #ab76493) was purchased from Abcam; The secondary antibodies including anti-rabbit immunoglobulin G (IgG) (H+L) (Dylight 800 conjugates, Cat. #5151) and anti-mouse IgG (H+L) (Dylight 680 conjugates, Cat. #5470) were also purchased from Cell Signaling Technology.

### Analyses of STING oligomerization and phosphorylation by TBK1 on liposomes

The liposomes were made by lipids 1-palmitoyl-2-oleoyl-sn-glycero-3-phosphoethanolamine (POPE, Avanti Research), 1-palmitoyl-2-oleoyl-glycero-3-phosphocholine (POPC, Avanti Research), brain L-α-phosphatidylserine (brain PS, Avanti Research), 1,2-dioleoyl-sn-glycero-3-phospho-(1’-myo-inositol-4’,5’-bisphosphate) (18:1 PI(4,5)P_2_, Avanti Research), 1,2-dioleoyl-sn-glycero-3-phospho-(1’-myo-inositol-3’,5’-bisphosphate) (18:1 PI(3,5)P_2_, Avanti Research), 1,2-dioleoyl-sn-glycero-3-phospho-(1’-myo-inositol-4’-phosphate) (18:1 PI(4)P, Avanti Research)) and cholesterol (Sigma-Aldrich). The lipids were dissolved in chloroform. The molar ratios of POPE and brain PS were fixed at 20% and 10%, respectively. Phosphatidylinositol-phosphates (PIPs) and cholesterol were added at defined molar ratios. The ratio of POPC was adjusted accordingly to ensure the total lipid composition summed to 100%. The lipid mixture was dried by nitrogen gas stream, followed by overnight vacuum to remove residual solvent. Dried lipids were hydrated and resuspended in buffer containing 20 mM HEPES (pH 7.4) and 150 mM NaCl. Lipid suspension was then extruded through a 0.2 µm polycarbonate membrane using an extruder, with 81 passes to produce uniformly sized liposomes. Liposomes were incubated with detergent n-decyl-β-D-maltoside (DM) at a final concentration of 0.5% (w/v) for 2.5 hours. Subsequently, full-length wild-type STING protein was added at a lipid-to-protein weight ratio of 20:1 and incubated for 1 hour. Incorporation of STING into the liposomes was achieved by gradual detergent removal via dialysis over 48 hours.

Cryo-EM micrographs of STING on liposomes of different lipid compositions were collected on the same Titan Krios microscopy as described above. The numbers of micrographs collected for the -PI(3,5)P_2_/-Cholesterol, -PI(3,5)P_2_/+Cholesterol, +PI(3,5)P_2_/-Cholesterol, +PI(3,5)P2/+Cholesterol conditions were 1204, 5211, 1971, and 2862, respectively. The length of STING filaments was measured in the program Napari v0.4 (https://napari.org/0.4.15/). The statistics were calculated using the Scipy statistics package v1.16 (https://docs.scipy.org/doc/scipy/reference/stats.html).

Recombinant human TBK1 (1-657) was expressed in sf9 cells and purified by His-tag affinity chromatography. STING in liposomes was incubated with 10 µM cGAMP for 15 minutes, followed by the addition of TBK1 and further incubation for 1 hour. The final concentration of STING and TBK1 were adjusted to 1 µM. Subsequently, ATP was added to a final concentration of 50 µM along with 5 mM MgCl₂ to initiate the reaction, which was carried out at 25 °C for 20 minutes. The reaction was terminated by adding SDS loading buffer, followed by heating at 80 °C for 3 minutes to denature the proteins. Phosphorylated and total STING protein levels were assessed by immunoblotting. The membranes were scanned with the Odyssey Infrared Imaging System (Li-COR, Lincoln, NE). Three or four independent experiments were performed. Data are mean ± s.d. P value was calculated with two-tailed Student’s t-test.

### Cell-based STING oligomerization assay

HEK293T cells were purchased from ATCC and maintained in Dulbecco’s modified Eagle’s medium (DMEM) supplemented with 10% fetal bovine serum (Sigma) and antibiotic antimycotic solution (Sigma). The coding sequence of full-length human STING followed by FLAG-tag was cloned into pcDNA3.1 (+) vector (Invitrogen). Mutations were introduced by Q5 mutagenesis kit. HEK293T expressing STING wild-type or mutants were treated with 1 μM cGAMP in the buffer containing 50 mM HEPES pH 7.5, 100 mM KCl, 3 mM MgCl_2_, 85 mM sucrose, 0.2% BSA, 1 mM ATP, 0.1 mM GTP and 10 μg/ml digitonin for 1 hour. Cells were resuspended in a buffer containing 20 mM HEPES pH 5.4, 150 mM NaCl, 1 mM DTT. Cells were lysed by sonication and debris was removed by centrifugation at 1,000 g for 5 minutes. The supernatant was incubated with 0.5% DDM and 0.05% CHS for 5 minutes and loaded onto native PAGE gels. Immunoblotting was performed with antibodies against STING.

### Cell-based phosphorylation assay

HEK293T cells were cultured in 6-welll plates and transfected with STING wild type and mutant plasmids. After being cultured for additional 20 hrs, the cells were stimulated with cGAMP at indicated concentration for 1 hr in a buffer containing 50 mM HEPES pH 7.5, 100 mM KCl, 3 mM MgCl_2_, 85 mM sucrose, 0.2% BSA, 1 mM ATP, 0.1 mM GTP and 10 μg/ml digitonin. Cells were incubated in cell lysis buffer (Cell Signaling Technology) and protease and phosphatase inhibitor (Thermo Scientific) for 20 minutes. After centrifugation at 21,130 g at 4 ℃, the concentration of cell lysate was measured by Micro BCA Protein Assay Kit (Thermo Scientific). Same amounts of proteins were loaded to SDS-PAGE and phosphorylation level of STING, TBK1 and IRF3 and α-tubulin was detected by immunoblotting. The membranes were scanned with the Odyssey Infrared Imaging System (Li-COR, Lincoln, NE).

### Immuno-fluorescence

The coding sequence of full-length human STING was cloned into the pmEGFP-N1 vector (Addgene). Mutations were introduced by Q5 mutagenesis kit. HEK293T cells were cultured on cover glasses in 6-well plates and transfected with STING wild type or mutant plasmids. After cells were cultured for additional 19 hrs, 1 μM cGAMP in the buffer containing 50 mM HEPES pH 7.5, 100 mM KCl, 3 mM MgCl_2_, 85 mM sucrose, 0.2% BSA, 1 mM ATP, 0.1 mM GTP and 5 μg/ml digitonin was used to stimulate cells for 1 hr. Cells were then washed by PBS and fixed with 4% paraformaldehyde. Nuclei were stained with 4’,6-diamidino-2-phenylindole (DAPI). Cells were imaged by a ZOE fluorescent cell imager (Bio-rad). Cells transfected with each STING construct were imaged in a blind manner in three independent experiments. The percentage of cells with STING-GFP puncta was calculated based on 150∼250 GFP-positive cells for each construct in each experiment. Images showing STING-GFP puncta formation were acquired on a Nikon Ti2E microscope equipped with a Yokogawa CSU X1 spinning disk using a 60× oil immersion objective with laser illumination and filter sets for FITC (488 nm laser) and DAPI (405 nm laser). Z-stacks were collected at 1 μm intervals through the cell volume. Image processing was done with FIJI^50^.

## Data availability

The coordinates and cryo-EM maps of the structures reported in the study have been deposited in the RCSB and EMD databases, respectively, with the following accession codes: 9DAN and EMD-46694 (STING/cGAMP/STG2/PI(4,5)P_2_), 9DAT and EMD-46697 (STING/cGAMP/STG2/PI(3,5)P_2_), 9DAV and EMD-46699 (STING/cGAMP/STG2/PI(4)P) and 9DAW and EMD-46700 (STING/cGAMP/C53/PI(4,5)P_2_)

## Acknowledgements

Cryo-EM data were collected at the University of Texas Southwestern Medical Center (UTSW) Cryo-Electron Microscopy Facility, funded in part by the Cancer Prevention and Research Institute of Texas (CPRIT) Core Facility Support Award RP220582. We thank D. Stoddard and J. Martinez Diaz for facility access. We thank the Structural Biology Laboratory at UTSW for equipment use. A portion of this research was supported by NIH grant U24GM129547 and performed at the PNCC at OHSU and accessed through EMSL (grid.436923.9), a DOE Office of Science User Facility sponsored by the Office of Biological and Environmental Research. We thank Marzia Miletto at PNCC for data collection. This work is supported in part by grants from the National Institutes of Health (R35GM150506 to J.X.T, R01-AI093967 to Z.J.C, R01CA273595 to X.Z. and X.-c.B., and R01CA299257 to Z.J.C., X.Z. and X.-c.B.), the Welch foundation (I-1389 to Z.J.C, I-1702 to X.Z. and I-1944 to X.-c.B.) and the Cancer Grand Challenge (CGCFUL-2021\100007) with support from Cancer Research UK and the US National Cancer Institute (to Z.J.C.). X.-c.B. and X.Z. are Virginia Murchison Linthicum Scholars in Medical Research at UTSW. Z.J.C. is a Howard Hughes Medical Institute (HHMI) investigator.

---

**Extended Data Fig. 1.**
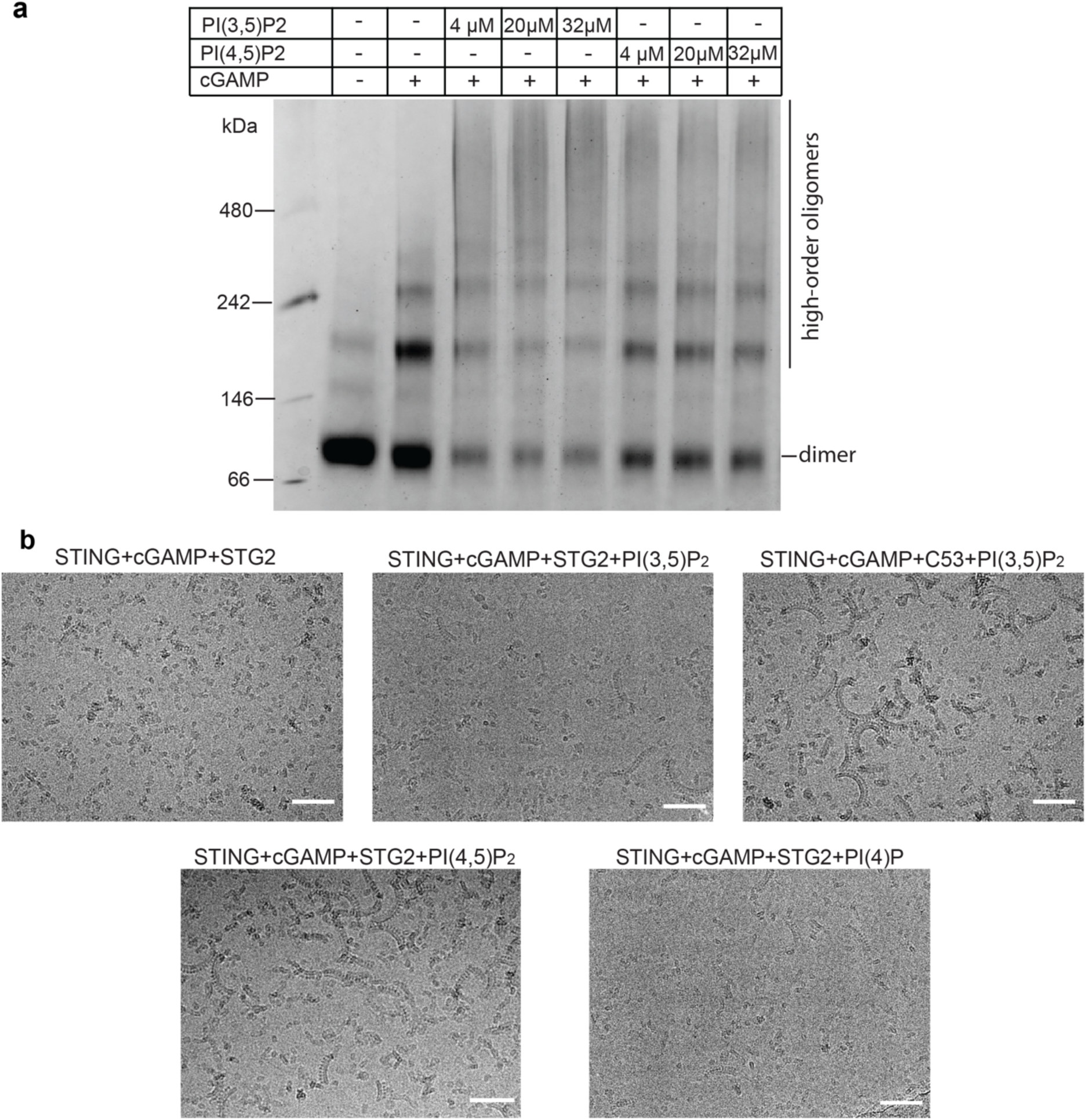
PIPs enhance the high-order oligomerization of STING to different extents. (**a**) Analyses of the high-order oligomerization of STING treated with various concentration of PI(3,5)P_2_ and PI(4,5)P_2_ by native PAGE. The image shown here is a representative of three biological repeats. (**b**) Cryo-EM micrographs showing high-order oligomers of STING bound to different combinations of agonists and PIPs. cGAMP and STG2 induced relatively short oligomers of STING. Different PIPs promoted the formation of longer oligomers to different extents. The micrographs are representative images from more than 5,000 distinct regions. The scale bar is 500 Å.

**Extended Data Fig. 2.**
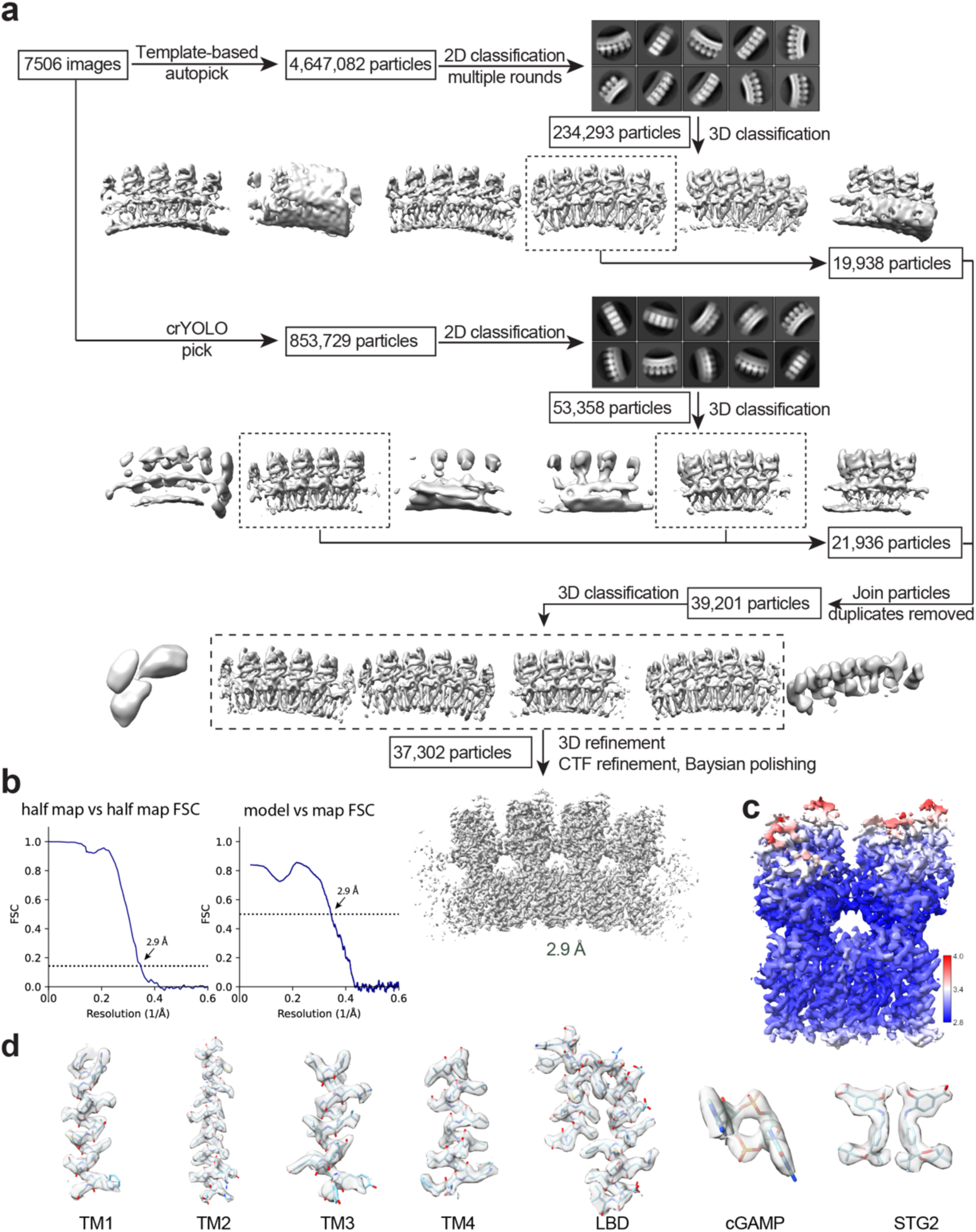
Image processing procedure for the cryo-EM structure of STING bound to cGAMP/PI(3,5)P_2_/STG2 and the quality of the structure. (**a**) Image processing procedure. (**b**) Gold-standard FSC curve of the final 3D reconstruction (left) and FSC between the map and atomic model (right). (**c**) Local resolution map of the final 3D reconstruction. (**d**) Sample densities of various parts of the structure.

**Extended Data Fig. 3.**
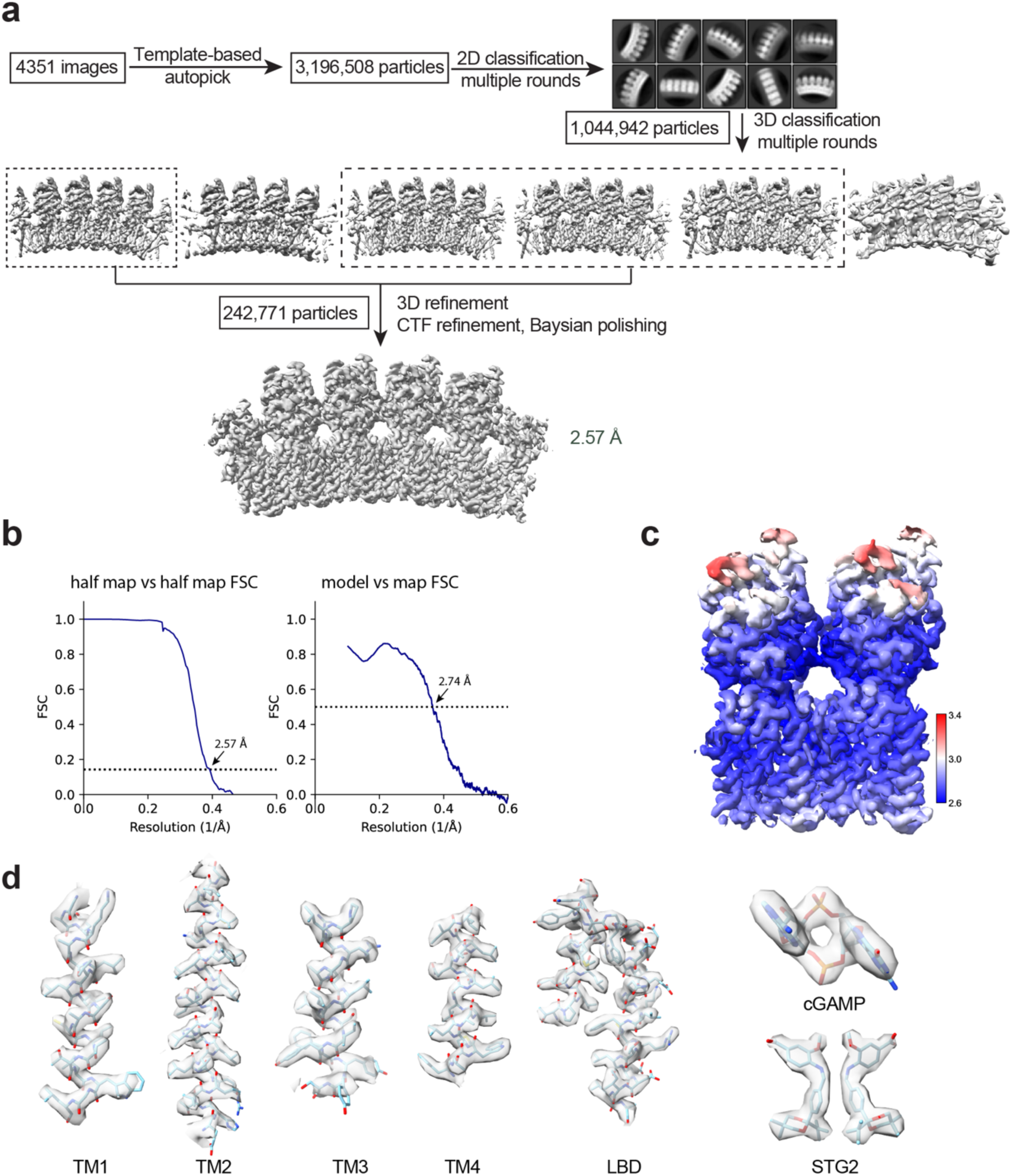
Image processing procedure for the cryo-EM structure of STING bound to cGAMP/PI(4,5)P_2_/STG2 and the quality of the structure. (**a**) Image processing procedure. (**b**) Gold-standard FSC curve of the final 3D reconstruction (left) and FSC between the map and atomic model (right). (**c**) Local resolution map of the final 3D reconstruction. (**d**) Sample densities of various parts of the structure.

**Extended Data Fig. 4.**
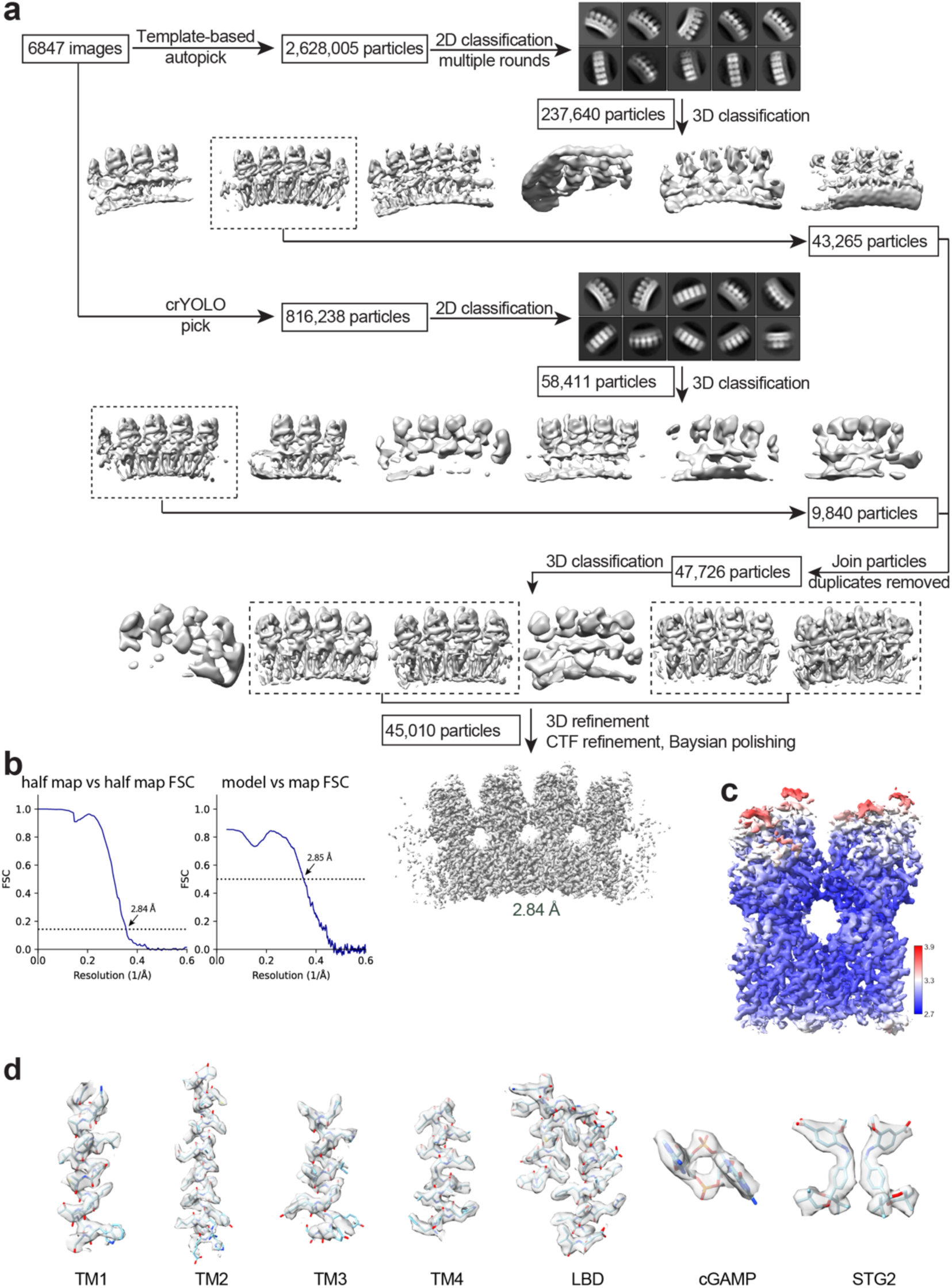
Image processing procedure for the cryo-EM structure of STING bound to cGAMP/PI(4)P/STG2 and the quality of the structure. (**a**) Image processing procedure. (**b**) Gold-standard FSC curve of the final 3D reconstruction (left) and FSC between the map and atomic model (right). (**c**) Local resolution map of the final 3D reconstruction. (**d**) Sample densities of various parts of the structure.

**Extended Data Fig. 5.**
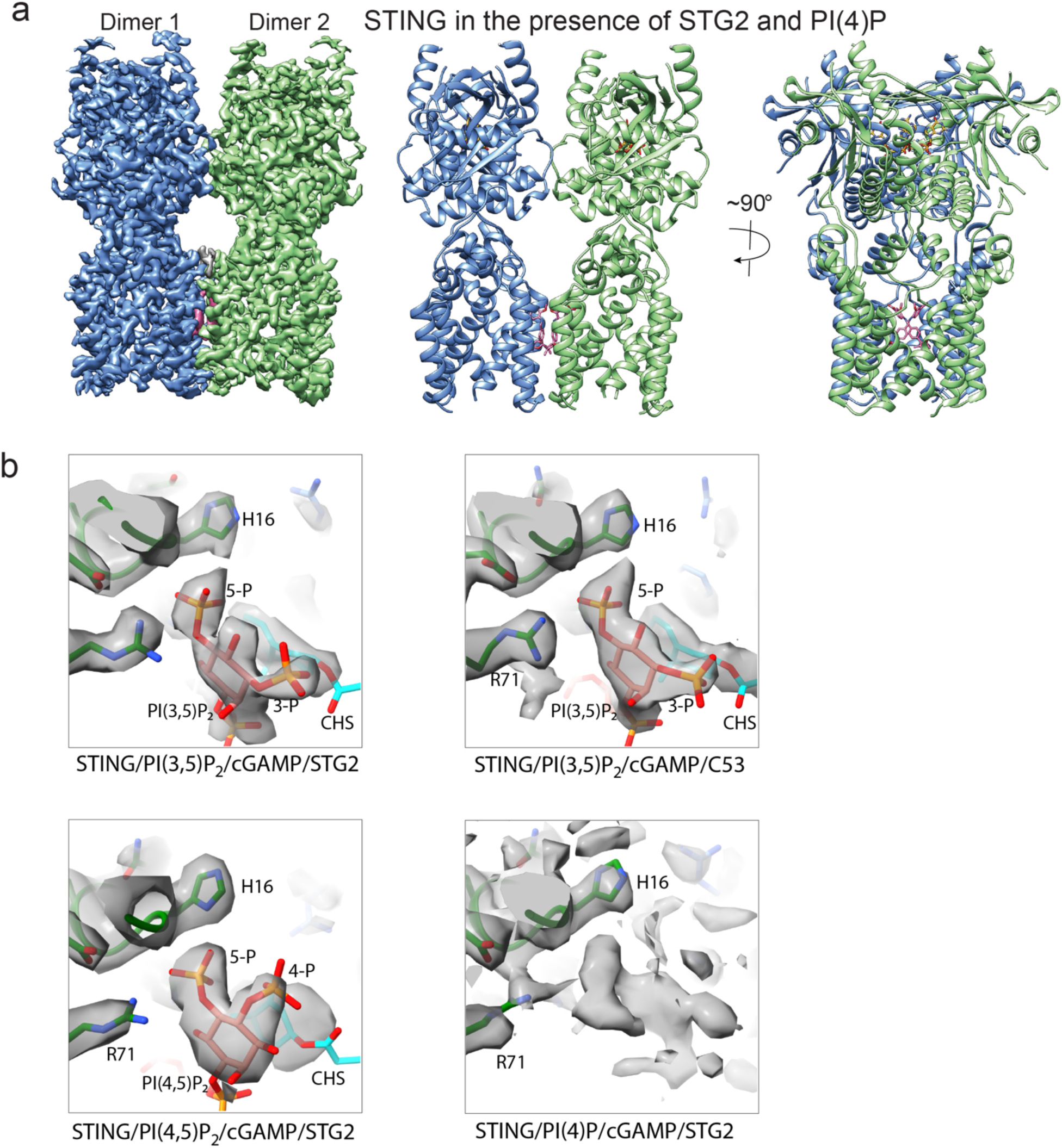
Comparison of the binding sites of different PIPs. (**a**) Cryo-EM map and atomic model of the STING high-order oligomer in the presence of cGAMP, PI(4)P and STG2. PI(4)P and CHS have very weak density and therefore not included in the atomic model. (**b**) Comparison of the density of PIPs in the four structures presented in this study. The thresholds of the cryo-EM maps were set such that the sidechain of H16 is covered at similar levels in the four structures. It is clear that the density for PI(4)P is much weaker than PI(3,5)P_2_ and PI(4,5)P_2_ in the three other structures.

**Extended Data Fig. 6.**
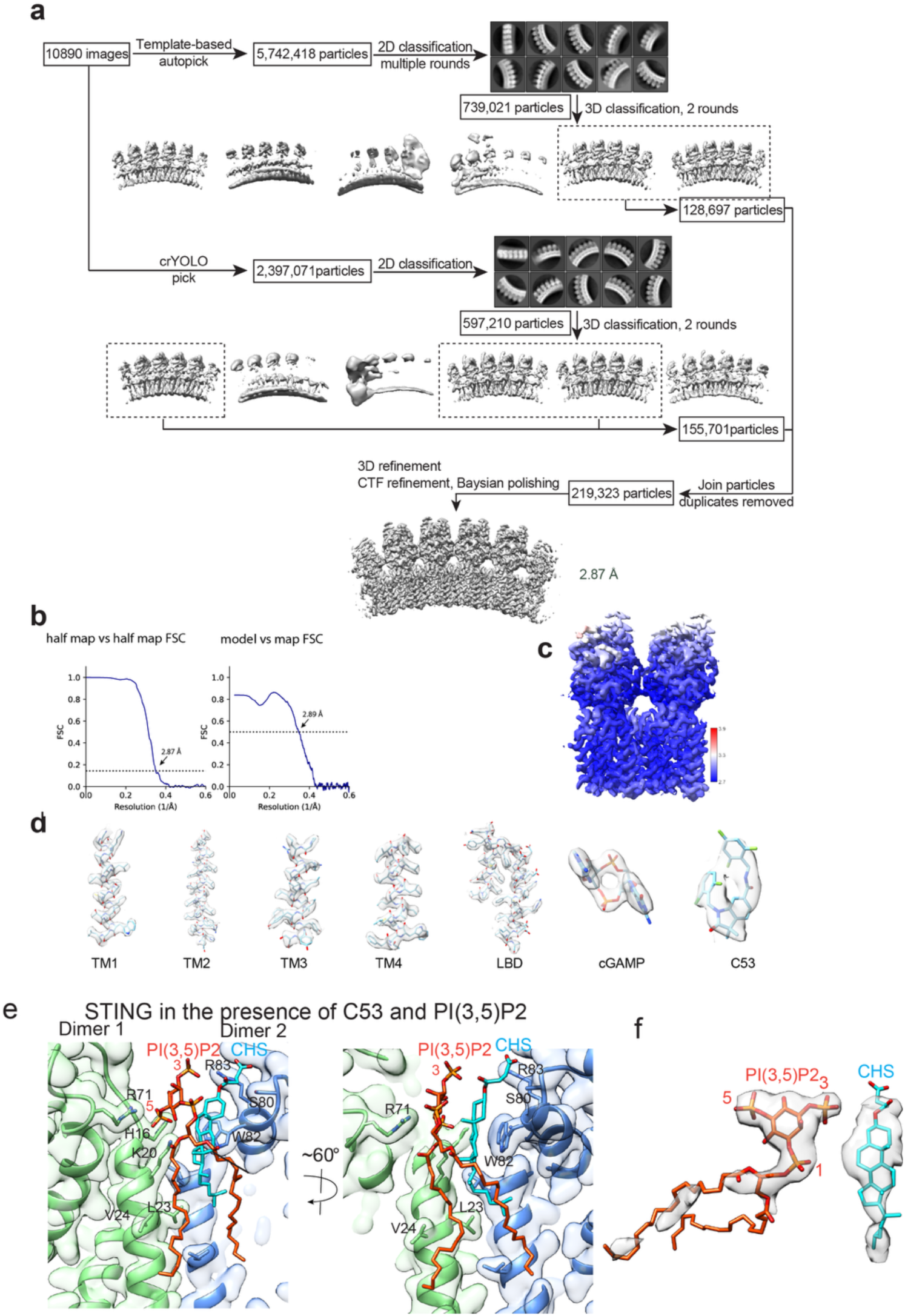
Image processing procedure for the cryo-EM structure of STING bound to cGAMP/PI(3,5)P_2_/C53 and the quality of the structure. (**a**) Image processing procedure. (**b**) Gold-standard FSC curve of the final 3D reconstruction (left) and FSC between the map and atomic model (right). (**c**) Local resolution map of the final 3D reconstruction. (**d**) Sample densities of various parts of the structure. (**e**) Detailed view of the binding modes of PI(3,5)P_2_ and CHS at the interface between two STING dimers in the structure of STING bound to cGAMP/PI(3,5)P_2_/C53. (**f**) Cryo-EM density of PI(3,5)P_2_ and CHS in the structure of STING bound to cGAMP/PI(3,5)P_2_/C53.

**Extended Data Fig. 7.**
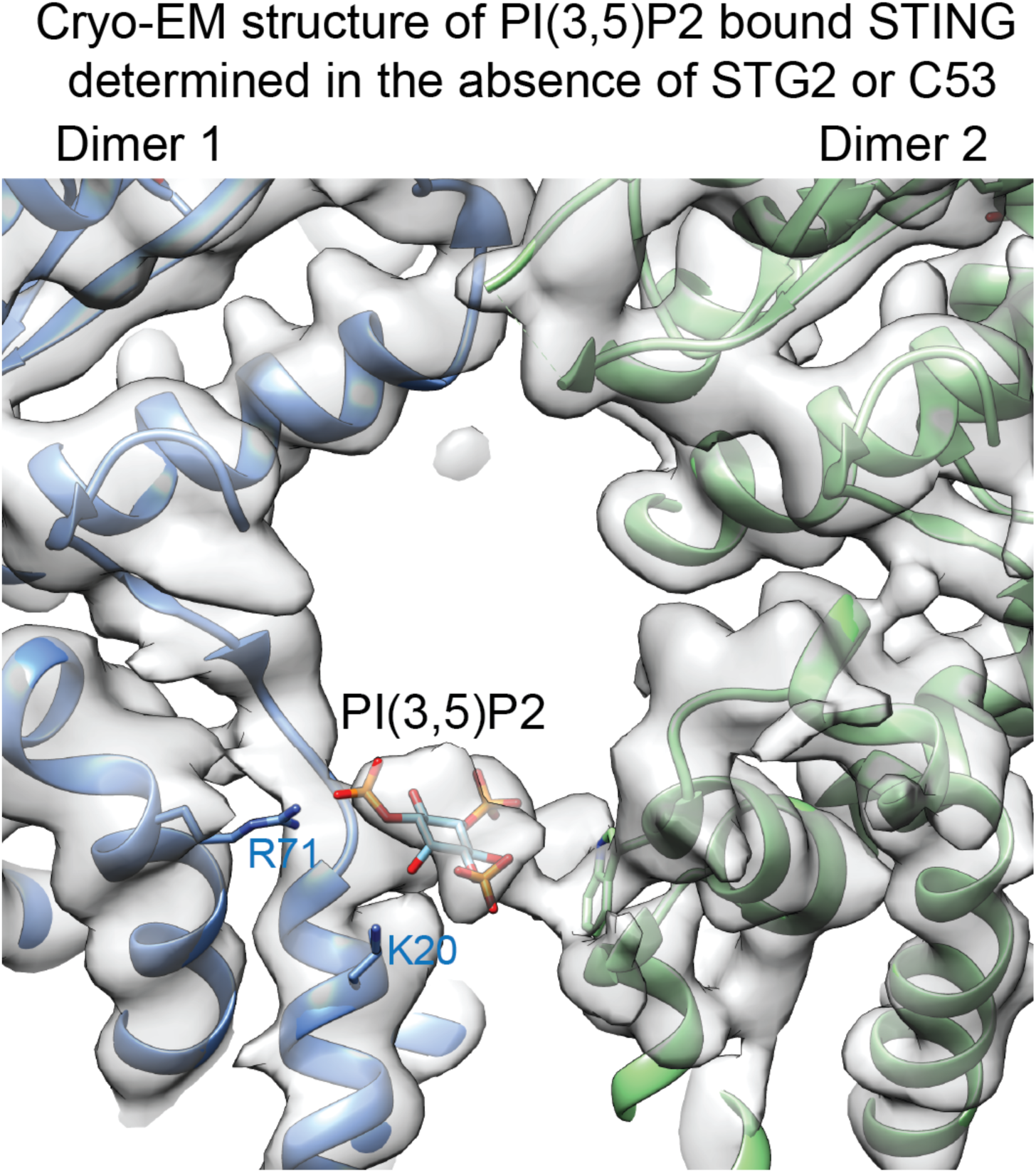
Cryo-EM map and atomic model of STING oligomer in complex with cGAMP and PI(3,5)P_2_ determined in the absence of STG2 or C53.

**Extended Data Fig. 8.**
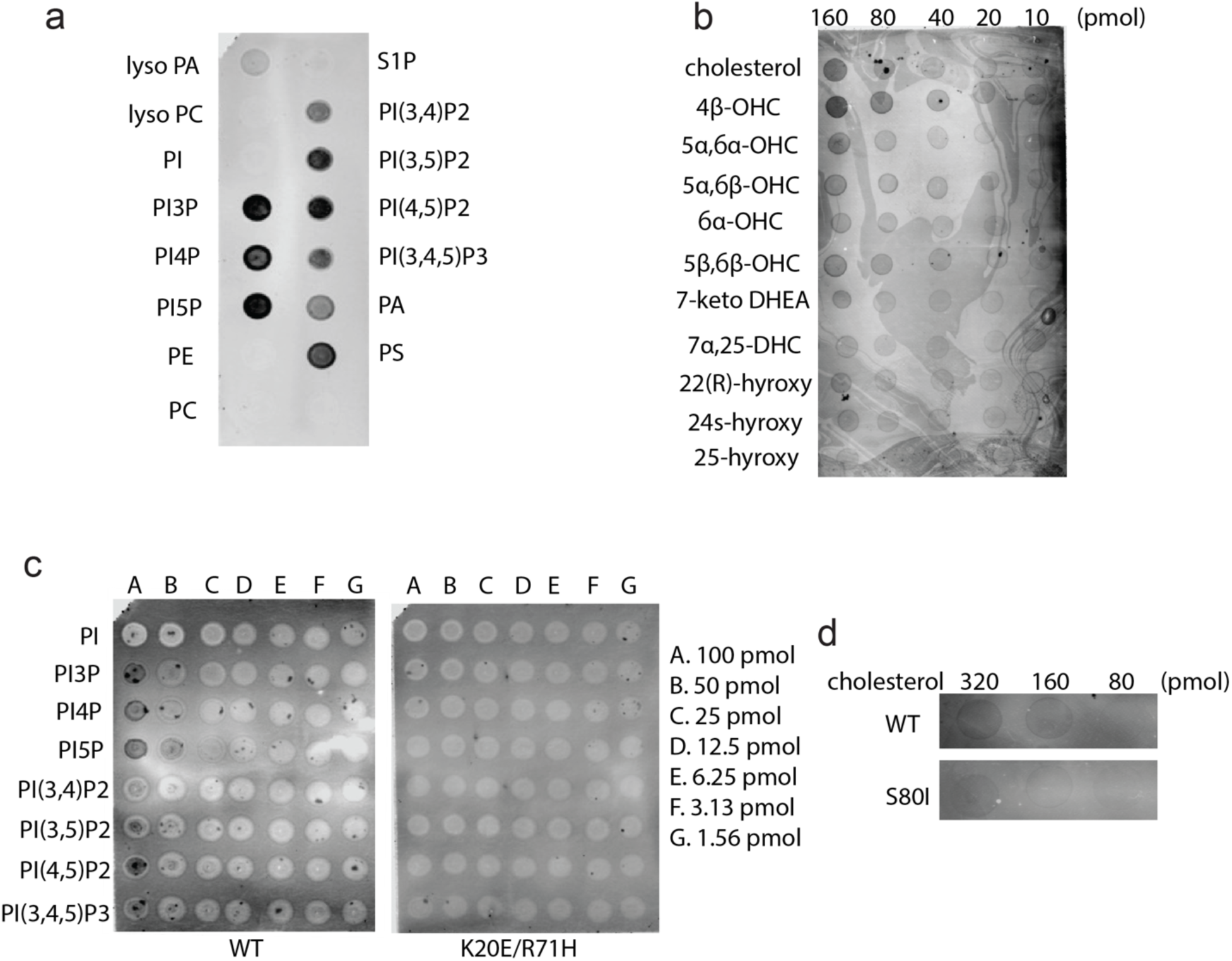
lipid binding assay for purified STING. (**a**) Binding of wild-type STING to a panel of lipids in a lipid-strip assay. (**b**) Binding of wild-type STING to an array of cholesterol and derivatives. (**c**) Comparison of binding to PIPs between wild-type STING and the K20E/R71H mutant. (**d**) Comparison of cholesterol binding between wild-type STING and the S80I mutant. The data shown are representative of three biological repeats.

**Extended Data Fig. 9.**
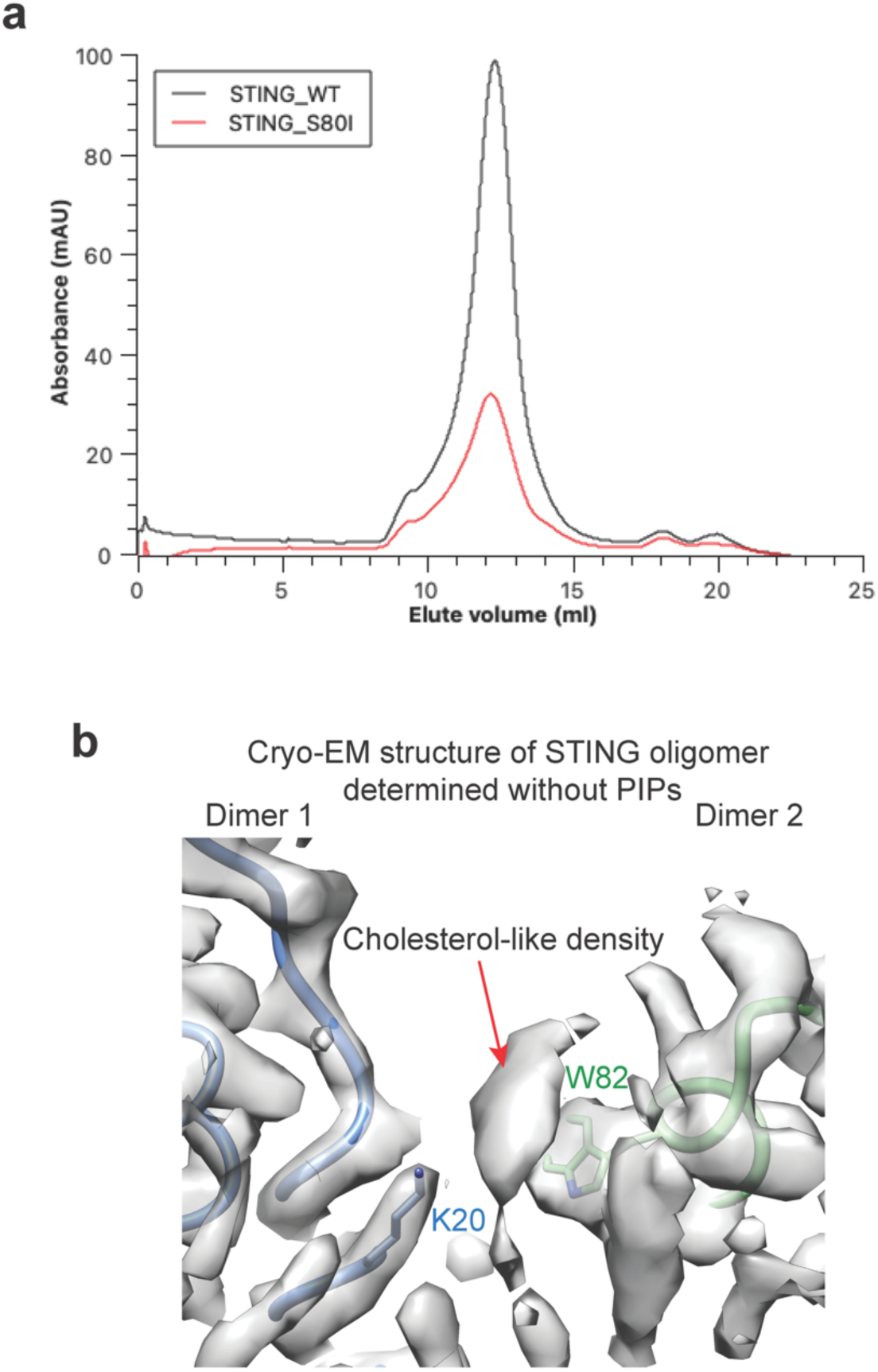
Cholesterol may bind to STING without PIPs. (**a**) The size exclusion chromatography profiles of wild-type STING and S80I mutant. The profile of the S80I mutant is similar to that of the wild type, suggesting that the S80I mutant is properly folded. (**b**) Cryo-EM map and atomic model of STING oligomer determined without PIPs (EMD-29282).

**Extended Data Table 1.**
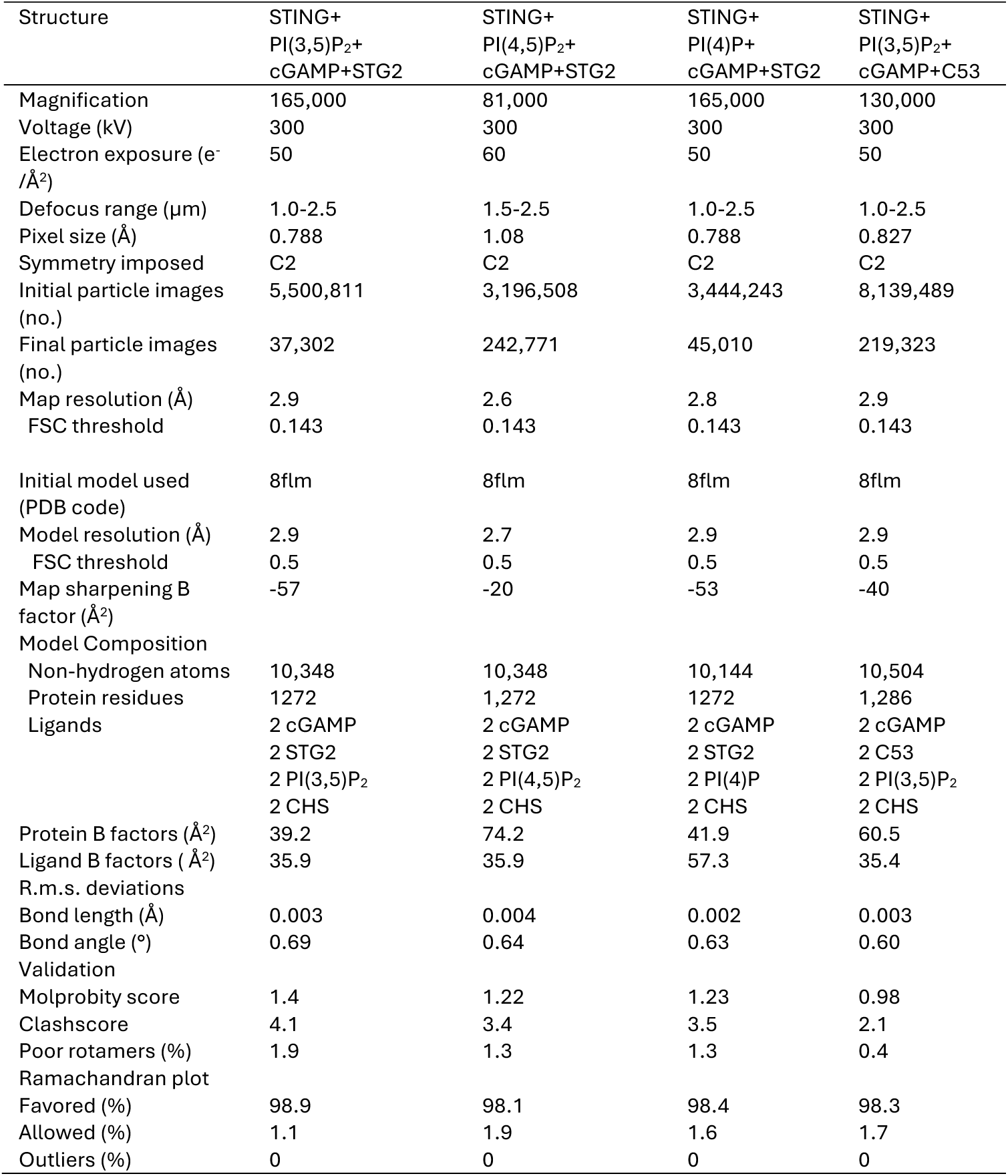
Cryo-EM data collection and structure refinement statistics.

